# Combination Treatment with Sclerostin and Dkk1 Antibodies Synergizes with Tibial Loading to Stimulate Bone Formation in Aged Mice

**DOI:** 10.64898/2026.02.23.707532

**Authors:** Lisa Y Lawson, Christopher J Chermside-Scabbo, Michael D Brodt, Nicole Migotsky, John T Shuster, Evan G Buettmann, Matthew J Silva

**Author notes:** **Corresponding Author:** Matthew J Silva, PhD, Department of Orthopaedic Surgery, Washington University, 425 S Euclid Ave, St. Louis, MO 63110, USA.

## Abstract

Aging is associated with decreased bone formation and bone mass and increased fracture risk. Wnt pathway activation by mechanical loading is a potent strategy to improve bone mass, however, load-induced bone formation is diminished with aging. Neutralizing antibody (Ab) therapies targeting Wnt pathway inhibitors Sclerostin (Scl) and Dickkopf-related protein 1 (Dkk1) have proven successful in preclinical and clinical osteoporotic conditions. We asked whether treatment combining Scl-Ab and Dkk1-Ab can increase load-induced bone formation in a preclinical model of osteoporosis. Aged (22-month) C57BL/6N female mice underwent combination Scl-Ab plus Dkk1-Ab therapy (15 mg/kg each; subcutaneously; 2x/wk; saline control) for 2 weeks, concomitant with a mechanical loading regimen previously shown to induce modest bone formation in tibias of aged mice (-2200 με, 1200 cycles/day, 5 day/wk). Changes in bone morphology and formation were assessed by longitudinal microCT and dynamic histomorphometry, respectively. Molecular indices of bone formation and Wnt pathway activation were assessed by qPCR of cortical bone. Treatment with Scl-Ab plus Dkk1-Ab induced significant improvements in cancellous (BV/TV +50%) and cortical morphology (Ct.Th +25%) in non-loaded limbs of antibody-treated mice vs. vehicle control mice. Importantly, periosteal bone formation rate was 10-fold higher in loaded limbs of antibody versus vehicle treated mice, indicating a synergistic effect. Gene expression analysis showed that antibody treatment and loading synergistically upregulated *Wnt1* expression, which may have contributed to the observed synergistic effect on bone formation. These results confirm the potent anabolic effect of combination Scl plus Dkk1 antibody treatment. Moreover, they show that antibody treatment and skeletal loading are more effective at increasing periosteal bone formation in aged mice than either treatment alone. These findings support the concept that combinatorial therapy using dual Scl and Dkk1 antibodies plus weight-bearing exercise may be an effective treatment for age-related osteoporosis.

## Introduction

Aging is associated with a progressive decline in bone mass and strength, leading to an increased risk of fractures. Bone mass and strength are maintained by a balance between bone resorption and formation. In this balance, the Wnt signaling pathway plays a critical regulatory role ^(1,2)^. Most early therapeutics for osteoporosis have worked by limiting bone resorption. However, these anti-catabolic treatments cannot reverse osteoporosis ^(3)^. Newer therapeutics, a class known as anabolic treatments, aim to stimulate bone formation to reverse low bone mass; some of these anabolic treatments work via modulation of Wnt signaling.

Two important extracellular regulators of the Wnt signaling pathway are the glycoproteins sclerostin (Scl, encoded by the gene *Sost*) ^(4,5)^ and Dickkopf-related protein 1 (Dkk1) ^(6,7)^. These secreted proteins inhibit pathway activation by binding to Wnt receptors LRP5/6, thus occupying the binding sites of endogenous activating Wnt ligands. Therefore, inhibiting Scl or Dkk1 via single neutralizing antibody treatment (monotherapy) should promote Wnt activation and downstream bone anabolism to treat osteoporosis. In support of this, the neutralizing Scl antibody (Scl-Ab) consistently increases bone formation and bone mass in animals ^(8,9)^ and humans ^(10,11)^ and has been FDA approved to treat post-menopausal osteoporosis. Yet, while Scl-Ab monotherapy leads to increased bone mass, it also increases *Dkk1* expression in bone, which may dampen its anabolic effect over time ^(12–14)^. Similarly, treatment with neutralizing Dkk1 antibody (Dkk1-Ab) increases sclerostin levels, which explains why Dkk1-Ab monotherapy does not reliably increase bone mass in wildtype adult rodents ^(15,16)^. However, when Dkk1-Ab treatment is given to *Sost*-null mice, it induces a robust increase in bone mass ^(16)^. Accordingly, there has been considerable interest in combination antibody therapy to neutralize both sclerostin and Dkk1, and studies have shown that dual therapy with Scl-Ab and Dkk1-Ab leads to greater increases in bone mass than with either antibody alone ^(13,16,17)^. More recently, bispecific antibodies that target both Scl and Dkk1 have shown efficacy in increasing bone mass in pre-clinical studies of adult mice ^(13)^ and are being tested in human clinical trials ^(18)^.

Another key regulator of the Wnt signaling pathway is mechanical loading ^(19–21)^, which is a well-established stimulus for new bone formation. Mechanical loading downregulates expression of the Wnt antagonists Scl and Dkk1 ^(22,23)^, and downregulation of *Sost* expression was shown to be required for the osteogenic response to loading ^(24)^. Moreover, loading upregulates expression of several Wnt ligands, including *Wnt1*, *Wnt7b*, *Wnt10b* and *Wnt16* ^(23,25yWergedal, 2015 #2621)^. The importance of Wnt ligands in the loading response was shown as blocking their secretion from osteoblasts significantly decreased load-induced bone formation ^(26)^. In particular, Wnt1 and Wnt16 have been shown to be necessary for loading-induced bone formation ^(27,28)^. Thus, mechanical loading potentiates Wnt signaling in bone in two ways, by downregulating Wnt antagonists and upregulating Wnt ligands, which are both necessary for maximal load-induced bone formation. Importantly, with aging the ability of loading to induce these Wnt pathway responses is blunted ^(23,29)^.

Given the links between Wnt signaling and mechanical loading in bone anabolism, several groups have investigated whether Wnt pathway neutralizing antibodies can boost the effects of mechanical loading. A synergistic effect of Scl-Ab treatment and loading was reported in young (2-month) ^(30)^ and young-adult (4-month) ^(31)^ mice. To date, it has not been reported whether a synergistic effect of Scl-Ab and loading is also observed in aged mice (which are a more clinically relevant model of osteoporosis), and more importantly, if combination Scl-Ab and Dkk1-Ab antibody treatment also elicits a synergistic effect when paired with loading. Thus, the goal of this study was to investigate the effect of dual Scl-Ab and Dkk1-Ab treatment on mechanical loading-induced bone formation in a preclinical model with relevance to osteoporosis. We hypothesized that dual Scl-Ab and Dkk1-Ab therapy in combination with mechanical loading increases bone formation compared to either loading or dual antibody treatment alone. To test this hypothesis, aged (22 month) female C57BL/6N mice were either given vehicle or combination Scl-Ab + Dkk1-Ab during a 2-week tibial loading regimen.

## Materials and Methods

### Mice

Studies were approved by the Washington University in St Louis IACUC. Adult 5-month female C57BL/6N wildtype mice (Charles River) were acquired and housed up to five per cage, given *ad libitum* access to chow and water, and aged to 22 months in a non-barrier facility on a 12/12 light/dark cycle. Mice were randomly assigned to one of two treatment groups: vehicle or combination Scl-Ab + Scl-Ab antibodies. All mice were subjected to axial tibial loading on the right limb. Sample sizes were selected to detect biologically meaningful differences in bone morphology and bone formation, and gene expression (n=10 mice/group, including allowance for sample attrition). All animals survived without adverse events. Mice were euthanized via CO_2_ asphyxiation.

### Drug Delivery

Dual antibody treatment was administered twice weekly by subcutaneous injection (Mon, Thurs), 1 hour before loading. Antibody-treated mice received a combination of Sclerostin (Scl-Ab) and Dkk1 (Dkk1-Ab) monoclonal antibodies (provided as a gift by Amgen) at a dose of 12.5 mg/kg each, as previously described ^(13)^. The antibody cocktail used for injections was prepared by diluting Scl-Ab and Dkk1-Ab to a final concentration of 3.6 mg/mL each in sterile PBS. Vehicle-treated controls were injected with 0.15 mL sterile PBS.

### In Vivo Mechanical Loading by Axial Tibial Compression

*In vivo* tibial loading was used to induce periosteal and endosteal lamellar bone formation, as described previously ^(23,32)^. With the mouse prone and anesthetized (1-3% isoflurane gas), the right leg (tibia) was placed vertically in the loading fixture. The knee was positioned superiorly in a semi-spherical cup (10 mm diameter) attached to the system actuator, and the foot was held in a static fixture inferiorly (20° dorsiflexion). A preload (−0.5 N) was applied, and the tibia was subjected to axial compression for 1200 cycles/day (4 Hz haversine waveform) using a materials testing system (Electropulse 1000, Instron). A peak force of -7 N was applied to engender a peak compressive strain of -2200 με, previously calibrated in aged 22-mo B6 female mice ^(33)^. This magnitude was selected with intent to induce a modest loading response in aged mice ^(23)^, allowing for additive or synergistic effects of antibody treatment to be detected. After each loading bout, buprenorphine (0.1 mg/kg subcutaneously) was delivered to mitigate pain from loading, and mice were returned to their cages to resume unrestricted activity. Contralateral, left tibias served as non-loaded controls. Mice were loaded either for 5 consecutive days for gene expression analysis (Fig. 1A), or 5 days/wk for 2 wks (10 days total) for bone morphometric and dynamic histomorphometry analyses (Fig. 2A).

**Figure 1.**
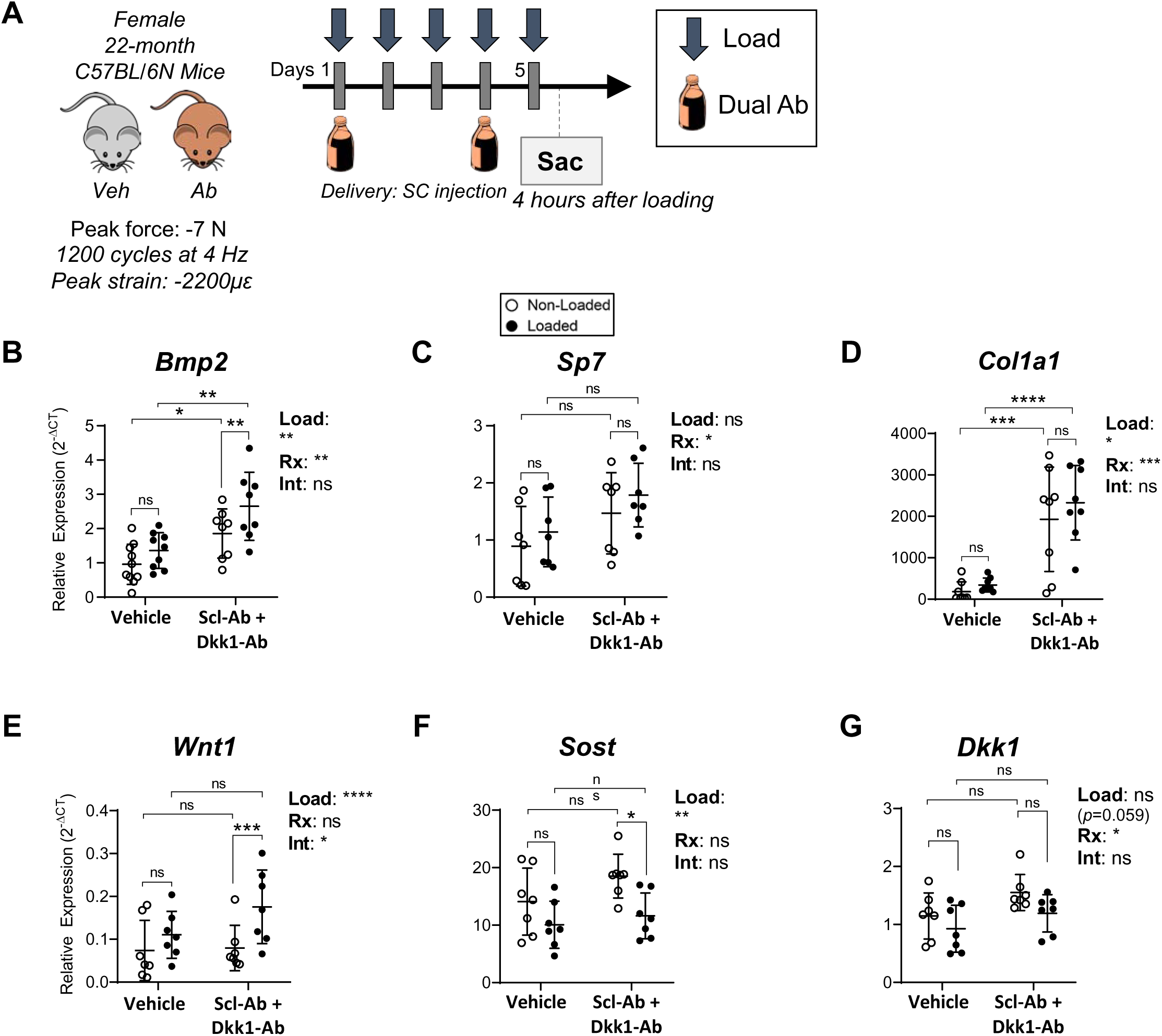
Antibody treatment and mechanical loading altered osteogenic and Wnt pathway gene expression. **(A)** Female 22-month-old mice were treated on days 1 and 4 with antibody (12.5 mg/kg Scl-Ab + 12.5 mg/kg Dkk1-Ab; subcutaneously) or vehicle, and loaded on days 1-5. Gene expression analysis was performed by RT-qPCR 4 hours following the last loading bout for osteogenic genes **(B)** *Bmp2*, **(C)** *Sp7*, and **(D)** *Col1a1*, and Wnt-related genes **(E)** *Wnt1*, **(F)** *Sost*, and **(G)** *Dkk1*. Two-factor repeated measures ANOVA was used to evaluate the main effects of loading (“Load”), dual antibody treatment (“Rx”), and their interaction. A significant loading-treatment interaction (“Int”) indicated that the transcriptional response to loading was differentially affected by treatment. Sidak test *post hoc* ANOVA was used for pairwise comparisons. (n=7-10 per group; 2-way repeated measures ANOVA with Šídák’s post-hoc test). All data shown as mean ± SD. * p<0.05, ** p<0.01, *** p<0.001, **** p<0.0001, ns p>0.05.

**Figure 2.**
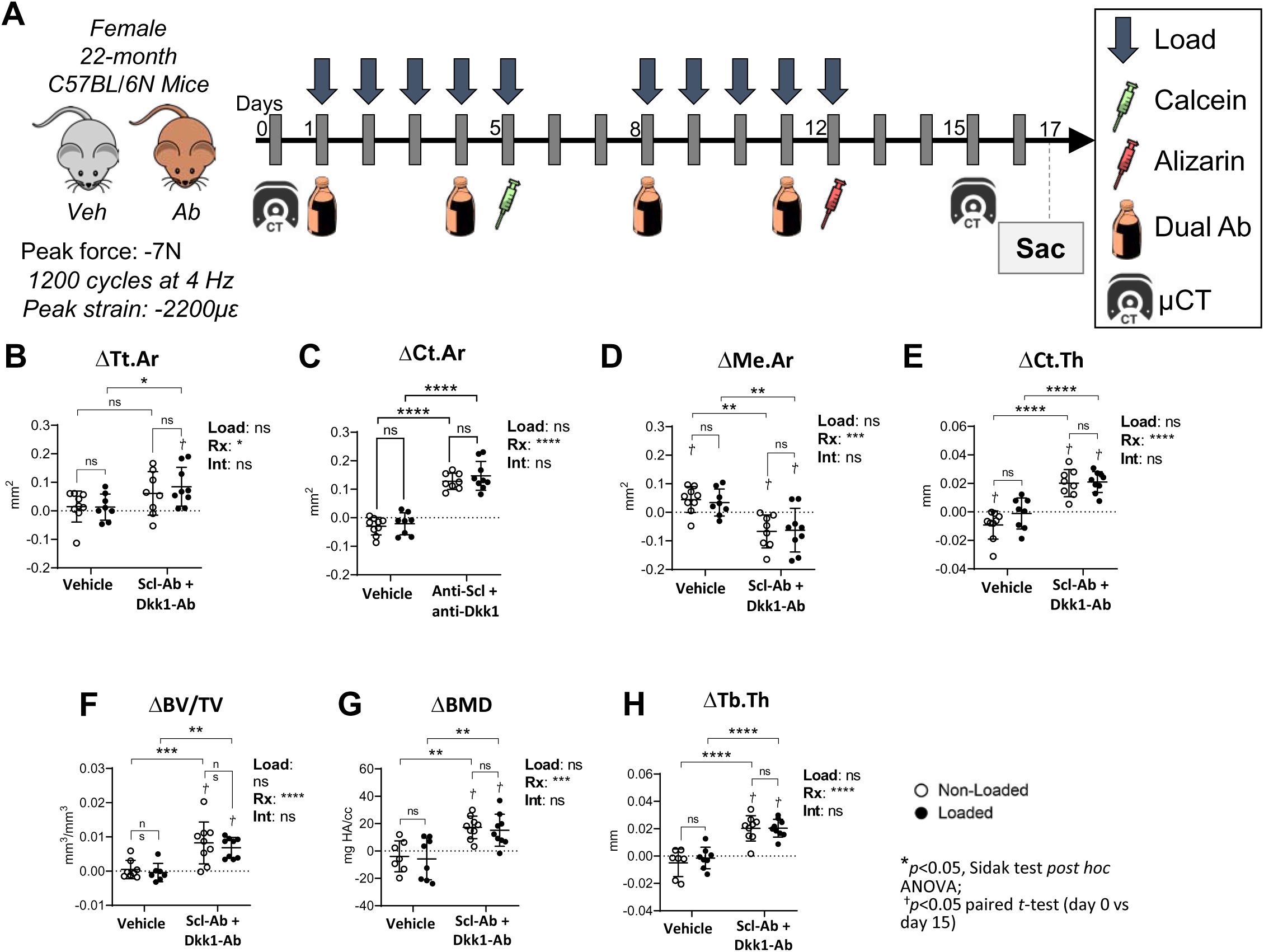
Antibody therapy enhances cortical and trabecular bone morphology. Serial µCT analysis was used to evaluate cortical (B-E) and trabecular (F-H) bone morphological changes (Δ) over a 2 wk treatment period. Two-factor repeated measures ANOVA was used to evaluate the main effect of loading (“Load”) and dual antibody treatment (“Rx”) on Δ (day 15 value *minus* day 0 value). Pairwise comparisons between groups were analyzed using Sidak’s multiple comparisons test *post hoc* ANOVA. \**p*<0.05, \*\**p*<0.01, \*\*\**p*<0.001, and \*\*\*\**p*<0.0001; n=8-9/group. D indicates that there was a significant change (*p*<0.05, paired *t*-test) in morphology between days 0 and 15 (refer to Figure S2A).

### Gene Expression

Genes related to osteoblast differentiation and Wnt pathway regulation were assayed by RT-qPCR. Tibias were harvested 4 hours after the final bout of loading following a 5-day loading protocol (Fig. 1A), as described previously ^(26)^. Briefly, bone marrow was removed from the medullary cavity, and the cortical diaphysis of each tibia was snap-frozen and pulverized for total RNA isolation. cDNA was prepared from samples with RNA integrity numbers (RINs) > 5.5 as assessed by Agilent 2100 Bioanalyzer. Gene expression was analyzed with SYBR-based reagents (StepOne Plus Machine, Applied Biosystems) or Taqman-based reagents (BioMark HD System) in collaboration with the Genome Technology Access Center at Washington University in St. Louis. Gene expression was normalized to reference gene *Tbp* and calculated using the 2^-ΔCT^ method.

### Serial µCT Analysis

*In vivo* serial µCT analysis was used to evaluate the effects of antibody treatment and loading on tibial morphology. Tibias were scanned on “day 0”, 1-2 days before initial treatment/loading and again on “day 15”, 3 days following the last bout of loading (Fig. 2A). Mice were anesthetized (1-3% isoflurane gas) and bilateral tibia were scanned at 21 µm voxel size, 70 kVp, 114 µA (VivaCT40, Scanco). A 2.1 mm region centered 5 mm proximal to the distal tibiofibular junction was used for cortical analysis, and a 1.5 mm region in the proximal metaphysis (immediately distal to the growth plate) was used for trabecular analysis. Standard microCT outcomes were determined using the manufacturer’s software ^(34)^. Changes in bone morphology from baseline were defined as Δ = value_Day 15_ – value_Day 0_. Mice were euthanized 2 days after the post-scan for bone formation analysis (described below).

### Dynamic Histomorphometry

In preparation for bone formation analysis by dynamic histomorphometry, mice were given calcein green (10 mg/kg, Sigma-Aldrich) and alizarin complexone (30 mg/kg, Sigma-Aldrich) by IP injection on days 5 and 12, respectively, and bilateral tibias were embedded in MMA (Fig. 2A). Thick (100 µm) transverse sections were cut ∼5 mm proximal to the distal tibiofibular junction and imaged on a confocal microscope (DMi8/TCS SPE, Leica Microsystems, Inc.). Calcein and alizarin-labeled surfaces were analyzed using Osteo II analysis software (Bioquant Image Analysis Corporation) to determine percent mineralizing surface (% MS/BS), mineral apposition rate (MAR), and bone formation rate (BFR/BS) on the periosteal (Ps) and endocortical (Ec) surfaces according to ASBMR standards ^(35)^. Relative bone formation rate (rBFR/BS), defined as BFR/BS_Loaded_ – BFR/BS_Non-Loaded_, was used as a measure of loading-induced bone formation.

### Statistical Analysis

Samples sizes (n=10/group) were chosen a priori based on historical effects sizes and variance from our previous loading studies using aged 22-month B6 mice and allowing for sample attrition ^(23,29)^. Final sample sizes ranged from 7-10 due to sample attrition for technical reasons. Statistical analysis was performed using Prism v7.0 (GraphPad Software, Inc.). Two-way ANOVA was used to evaluate the main effects of loading (“Load”: loaded vs. non-loaded control), treatment (“Rx”: Scl-Ab+Dkk1-Ab vs. vehicle control), and the loading- Rx treatment interaction (“Int”), followed by Sidak’s multiple comparisons test for pairwise comparisons. One-factor ANOVA was used to compare relative bone formation rates (rBFR/BS) in Ab vs vehicle-treated mice. Paired *t*-tests were used to compare bone morphology from µCT analysis in non-loaded limbs on day 15 vs day 0. Significance was defined as \**p*<0.05, \*\**p*<0.01, \*\*\**p*<0.001, and \*\*\*\**p*<0.0001, with trends noted at 0.05 < *p* < 0.10. Individual data points and the mean ± standard deviation were plotted in each graph; each data point represents one animal; sample sizes in figure captions.

## Results

### Dual antibody therapy and loading additively upregulated osteogenic gene expression

To evaluate the combined effects of dual Scl-Ab and Dkk1-Ab therapy and mechanical loading on osteogenic and Wnt pathway-related gene expression, qPCR analysis was performed on cortical bone from bilateral tibias after 5 days of loading. Loading and antibody treatment each significantly increased the expression of the osteogenic genes *Bmp2* and *Col1a1* in the tibia (main effects in 2-factor ANOVA, p<0.05; Figs. 1B, 1D). There was no significant interaction (“int” effect in 2-factor ANOVA) between loading and treatment on *Bmp2* and *Col1a1* expression, so their net effects were additive. Antibody treatment (but not loading) also significantly upregulated osterix (*Sp7*) expression (Fig. 1C). Neither loading nor antibody treatment significantly affected the expression of osteocalcin (*Bglap)*, *Runx2*, or *Dmp1* (Fig. S1A-C). Because there were no significant interactions between loading and treatment, their effects on osteogenic gene expression appear to be independent and additive.

### Loading, but not dual antibody therapy, induced upregulation of Wnt ligands

A survey of mechanosensitive Wnt pathway-related genes showed that loading increased expression of the ligands *Wnt1*, *Wnt7b*, and *Wnt16* (loading main effect, *p*<0.05), whereas antibody treatment did not (Figs. 1E and S1E-F). Loading (main effect) also had a significant or near-significant inhibitory effect on *Sost* and *Dkk1* expression, respectively (Fig. 1F-G). Two-factor ANOVA indicated a modest but significant elevation in *Dkk1* expression in the bones of antibody-treated mice (Rx main effect, p<0.05; Fig. 1G). Neither loading nor treatment significantly affected the expression of canonical Wnt target gene *Axin2* (Fig. S1D). Finally, expression of *Wnt1* showed a significant loading-treatment interaction (Fig. 1E), suggesting that antibody treatment potentiated the effect of loading. However, no other Wnt-pathway related genes demonstrated a loading-antibody interaction. In sum, loading and antibody-mediated neutralization of Scl and Dkk1 independently increased osteogenic gene expression in aged mice, while only loading consistently altered expression of Wnt-pathway related genes, with a mild but significant interactive effect on *Wnt1* expression.

### Dual antibody therapy, but not loading, significantly enhanced cortical and trabecular bone morphology

Serial μCT analysis was used to evaluate the effects of antibody treatment and loading on bone morphometric changes over a 15-day period. Mice were treated with antibodies twice weekly and subjected to daily loading for 2 weeks (Fig. 2A). Left (non-loaded) and right (loaded) tibias were pre-scanned (day 0) and post-scanned (day 15) to calculate change in bone morphology from baseline (Δ = day 15 value minus day 0 value).

Dual antibody treatment had a significant effect on bone morphology, independent of loading (Figs. 2B-H and S2B). In non-loaded limbs of antibody-treated mice, cortical area (Ct.Ar) increased by 24% (day 15 vs day 0, *p*<0.0001; Fig. S2A). Similarly, in non-loaded limbs from antibody-treated mice cortical thickness (Ct.Th) and moment of inertia (pMOI) increased by 15% and 24%, respectively (*p*<0.01; Fig. S2A). In contrast, in non-loaded limbs from vehicle-treated controls, Ct.Ar, Ct.Th, and pMOI either remained unchanged or decreased slightly (-5% to -9%; *p*<0.05, paired *t*-test) between day 0 and day 15 (Fig. S2A). Additionally, medullary area (Me.Ar) increased slightly in the non-loaded limbs of vehicle-treated controls over the 15-day period (+5%, *p*<0.05, paired *t*-test), whereas in antibody-treated mice Me.Ar decreased (-9%, *p*<0.05, paired *t*-test), suggesting that antibody treatment protected against endocortical bone loss (Fig. S2A). In contrast, tibial loading did not significantly affect changes in measured cortical bone morphology that occurred during the 15-day study period, as evidenced by no significant main effects of loading on cortical parameters ΔTt.Ar, ΔCt.Ar, ΔMe.Ar, or ΔCt.Th (Fig. 2B-E).

Dual antibody treatment had a similar positive effect on trabecular bone morphology, also independent of loading. For example, antibody treatment (but not loading) increased BV/TV, BMD, and Tb.Th (Fig. 2F-H). In non-loaded limbs from antibody-treated mice, BV/TV was 50% greater on day 15 vs day 0 (*p*<0.01; Fig. S2A), while Tb.Th and BMD were 31-34% greater on day 15 vs day 0 (*p*<0.001; Fig. S2A). By comparison, in vehicle-treated mice BV/TV, BMD, and Tb.Th were not different on day 15 vs day 0 in either limb (*p*>0.05, paired *t*-test) (Fig. S2A). In summary, analysis of temporal changes in bone morphology indicated that antibody treatment led to significant increases in measures of cortical and trabecular bone quantity, whereas loading did not significantly alter bone morphology during the 15-day study period.

### Dual antibody therapy and loading synergistically increased periosteal bone formation

Bilateral tibias were harvested on day 17 and processed for analysis of diaphyseal bone formation by classic dynamic histomorphometry (Fig. 3A). Both loading and antibody treatments significantly increased measures of periosteal bone formation. Two-factor ANOVA indicated that loading increased periosteal mineralizing surface (Ps.MS/BS), mineral apposition rate (Ps.MAR) and bone formation rate (Ps.BFR/BS) (“Load” main effect, *p*<0.01; Fig. 3B). Pairwise comparisons showed that in mice treated with Scl-Ab plus Dkk1-Ab, loading increased Ps.MS/BS by 1.9-fold (loaded vs non-loaded, *p*<0.0001), Ps.MAR by 1.8-fold (*p*<0.01), and Ps.BFR/BS by 3.7-fold (*p*<0.0001). In contrast, loading had negligible effects in vehicle-treated controls; pairwise comparisons showed no difference in periosteal parameters between loaded vs non-loaded limbs of vehicle-treated mice (Fig. 3B; p>0.05 by Sidak post-hoc).

**Figure 3.**
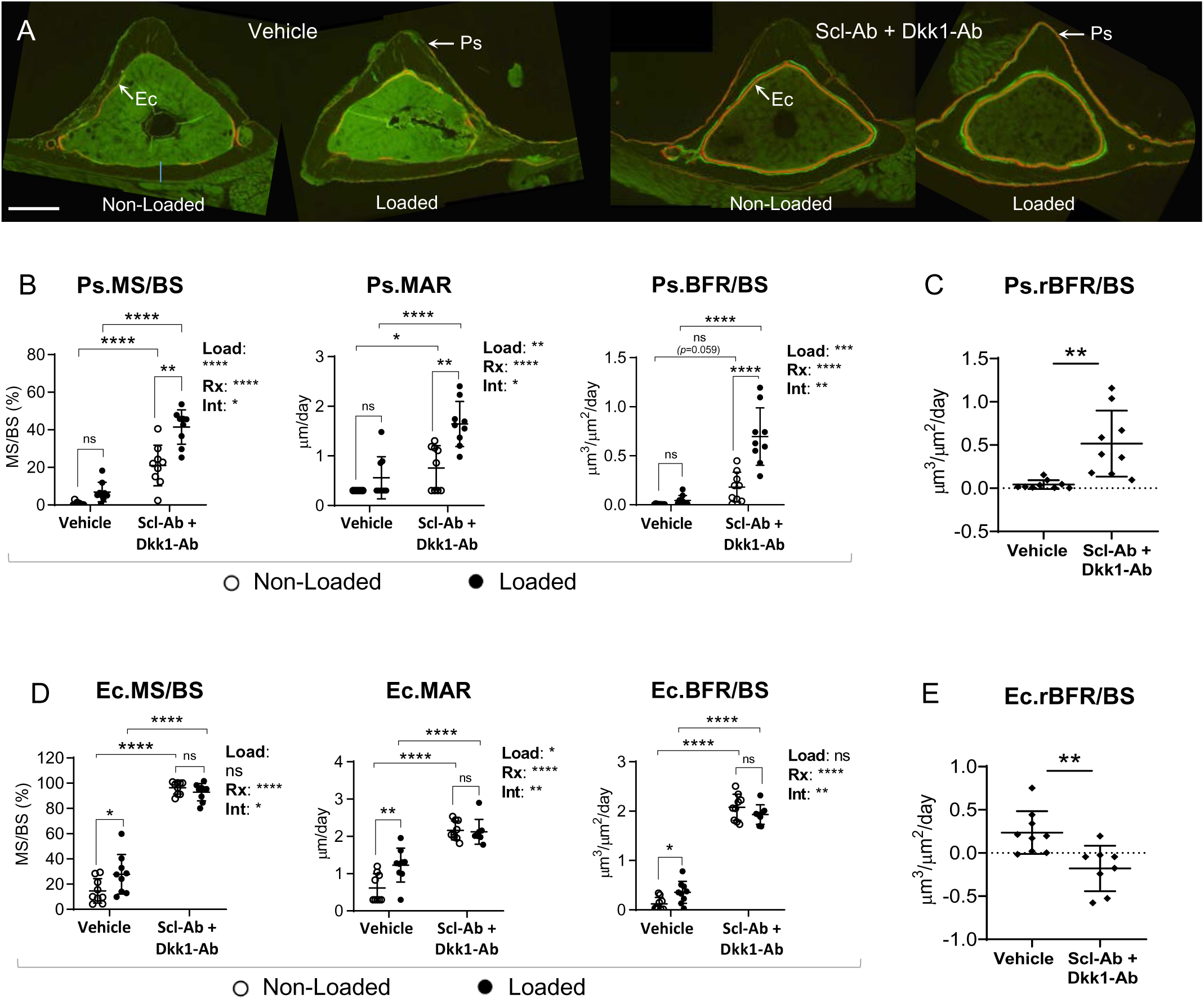
Loading and antibody therapy act synergistically to increase periosteal bone formation. Bone formation was evaluated on day 17, following calcein green and alizarin red injections on days 5 and 12, respectively. (A) Representative images showing fluorophore label on the endocortical (Ec) and periosteal (Ps) bone surface in each group. Scale bar=250 µm. (B, D) Two-factor ANOVA was used to evaluate the effects of loading (“Load”), dual antibody treatment (“Rx”), and their interaction (“Int”), followed by Sidak test for pairwise comparisons. (C, E) Relative bone formation rate (rBFR/BS), defined as BFR/BS_Loaded_ minus BFR/BS_Non-Loaded_, was used as an overall measure of the bone anabolic response to loading. \**p*<0.05; n=9/group.

Dual antibody therapy also significantly increased periosteal bone formation in both non-loaded and loaded limbs (“Rx” main effect, *p*<0.0001; Fig. 3B). Specifically, in non-loaded limbs antibody therapy increased Ps.MS/BS (35-fold; 21 vs 0.6%, Ab vs Veh groups, *p*<0.0001), Ps.MAR (2.5-fold; 0.76 vs 0.30 µm/day, *p*<0.05), and Ps.BFR/BS (90-fold; 0.18 vs 0.002 µm^3^/µm^2^/day, *p*=0.059).

Importantly, dual antibody treatment significantly enhanced the periosteal response to loading, demonstrated by significant (or near-significant) interaction effects by two-way ANOVA (“Int”, *p*<0.05 for Ps.MS/BS and Ps.BFR/BS, *p*=0.054 for Ps.MAR; Fig. 3B). Specifically, pair-wise comparison of loaded limbs mice showed that Ps.MS/BS and Ps.MAR were 4.6- and 2.2-fold higher, respectively, in antibody versus vehicle treated mice while Ps.BFR/BS was 9.8-fold higher in loaded limbs of antibody versus vehicle treated mice (*p*<0.0001; Fig. 3B). The net effect of loading on bone formation in each group was also analyzed by calculating the relative bone formation rate (rBFR/BS = loaded minus non-loaded) for each mouse. Compared to vehicle-treated controls, aged mice treated with Scl-Ab + Dkk1-Ab had a significantly greater relative bone formation response to tibial loading (12.3-fold; ANOVA, *p*<0.01; Fig. 3C). In summary, dual antibody therapy significantly enhances the anabolic effects of mechanical loading on the periosteal surface, indicating a synergistic effect of the two interventions.

### Dual antibody therapy, independent of loading, potently induced endocortical bone formation

Endocortical bone formation was robustly increased by Scl-Ab + Dkk1-Ab antibody treatment with average values of tibial Ec.MS/BS approaching 100% (Fig. 3A,D). By contrast, the effect of loading on the endocortical surface was minimal and observed only in vehicle-treated mice. Two-factor ANOVA indicated that antibody treatment had a strong and highly significant effect on Ec.MS/BS, Ec.MAR, and Ec.BFR/BS (“Rx” main effect, *p*<0.0001; Fig. 3D). Pairwise comparisons of non-loaded bones from vehicle vs antibody-treated mice showed that Ec.MS/BS was 6.6-fold higher in antibody-treated mice (*p*<0.0001), Ec.MAR was 3.5-fold higher (*p*<0.0001), and Ec.BFR/BS was 17.3-fold higher (p<0.0001; Fig. 3D). Pairwise comparisons of loaded bones from vehicle vs antibody-treated mice showed similar trends, with Ec.MS/BS, Ec.MAR, and Ec.BFR/BS being 3.3-fold, 1.7-fold, and 9.8-fold higher in antibody-treated mice (p<0.0001; Fig. 3D).

Unlike in the periosteum, dual antibody treatment did not further enhance the endocortical response to tibial loading. Although there was a significant interaction between antibody treatment and loading (“Int”, *p*<0.05 for Ec.MS/BS, Ec.MAR, and Ec.BFR/BS), this was because loading significantly enhanced endocortical bone formation in vehicle-treated mice but not in antibody-treated mice. For example, in vehicle-treated mice, values of Ec.BFR/BS were 3-fold higher in loaded limbs vs non-loaded limbs (p <0.05, Fig. 3D). In contrast, comparisons between loaded and non-loaded bones in antibody-treated mice showed no significant loading effect (p>0.05; Sidak post-hoc test; Fig. 3D). Consequently, the relative endosteal bone formation rate (rBFR/BS) showed a modestly diminished bone formation response to tibial loading in antibody versus vehicle-treated mice (*p*<0.05; Fig. 3E). While this result may suggest a suppressive effect of antibody treatment on the endocortical bone formation response to loading, it seems more likely that dual antibody treatment alone was so potent (average Ec.MS/BS approaching 100%) that loading did not further stimulate bone formation on this surface.

In sum, these findings show that both mechanical loading and dual antibody therapy improved bone formation outcomes in tibias of aged mice, albeit to differing extents on the periosteal versus endosteal surfaces. On the periosteal surface, antibody treatment and loading synergistically increased bone formation while on the endosteal surface, antibody treatment alone robustly increased bone formation so much as to mask any effect of additional mechanical loading.

## Discussion

We sought to test the hypothesis that pharmacological activation of the Wnt pathway by dual Scl-Ab + Dkk1-Ab treatment enhances loading-induced bone formation in aged mice. Our results support the hypothesis, as we observed a synergistic effect of loading and antibody treatment on periosteal bone formation; Ps.BFR/BS was 10-fold greater in loaded tibias from antibody-treated mice compared to vehicle-treated mice. On the endosteal surface, antibody treatment alone resulted in ∼100% mineralizing surface with no additional benefit of mechanical loading based on dynamic histomorphometry. In addition, despite antibody treatment by itself significantly increasing micro-CT measures of cancellous and cortical bone structure, we observed no significant additive or synergistic effect of loading on these outcomes in this 2-week study. Lastly, we did observe a synergistic effect of loading and dual antibody treatment on *Wnt1* expression, and an additive effect on expression of *Bmp2* and *Col1a1*. In all, our results indicate that loading can enhance the bone anabolic effects of dual Scl + Dkk1 antibody therapy in the aged murine skeleton, supporting the concept that antibody treatment and weight-bearing exercise may be a beneficial combination in osteoporotic individuals.

### Mechanical loading and Scl+Dkk1 antibody treatment synergistically increase periosteal bone formation in aged mice

We saw no differences between loaded and non-loaded tibias of vehicle-treated animals in periosteal dynamic histomorphometry (Fig. 3B,C), or in cortical or cancellous bone morphology by microCT (Fig. 2B-H). These findings indicate that the loading regimen we used was insufficient to induce a significant bone formation response in aged C57Bl/6 female mice; this is in slight contrast with our prior results showing that an equivalent loading regimen induced a modest bone formation response in mice, albeit one that was significantly less than in young-adult mice ^(29,36)^. Regardless, the lack of a loading response in vehicle-treated mice in the current study provided a suitable condition to test our hypothesis. In antibody-treated mice, loading induced a significant increase in periosteal bone formation, indicating a synergistic, anabolic effect of the two factors (Fig. 3B,C). These results build on previous findings on the combined effects of Scl-Ab with loading in young and ovariectomized mice. Morse et al. and Gerbaix et al. ^(30,31)^ reported that mechanical loading and Scl-Ab synergistically increased measures of bone formation and bone structure in young-adult mice. The present study extends these results to the aged, osteoporotic skeleton and demonstrate that the impaired responsiveness to loading with age can be enhanced by dual Scl + Dkk1 antibody therapy.

### Synergistic effects of load and antibody treatment on periosteal bone formation may involve Wnt1

Expression of Wnts 1, 7b and 16 were significantly upregulated by mechanical loading but not antibody treatment (Fig. 1E, S1 E-F). In addition, expression of WNT antagonists *Sost* and *Dkk1* were downregulated by loading (Fig. 1F,G), with only *Dkk1* expression having a modest upregulation with antibody treatment. Thus, mechanical loading, even in the aged skeleton, upregulated Wnt ligands and downregulated Wnt antagonists, effects that are known to contribute to a bone anabolic response ^(24,27,28)^. This may explain why loading enhanced the effect of antibody treatment, which had little to no effect on expression of these genes. Notably, *Wnt1* was the only gene measured whose expression showed a significant interaction between mechanical loading and dual antibody therapy (Fig. 1E), a response that mirrored the synergistic periosteal bone formation response (Fig. 3B). This suggests that a mechanism by which Scl-Ab + Dkk1-Ab synergistically increased periosteal bone formation in the current study was via enhancing load-mediated *Wnt1* upregulation in aged bone. This synergistic molecular effect is in line with the findings of Morse et al. ^(30)^, who showed that loading and Scl-Ab treatment in young mice synergistically enhanced WNT/β-catenin and Rho GTPase pathways compared to either treatment alone; each of these pathways are downstream of Wnt activation in bone^(2)^. However, the mechanism linking dual antibody treatment with synergistic upregulation of *Wnt1* gene expression with loading, as we observed, is unclear. It is possible that the mechanosensitive Piezo1/YAP/TAZ axis is involved. Piezo1 directly regulates differential *Wnt1* and *Sost* expression in loaded osteocytes in part via YAP/TAZ, and is necessary for load induced bone formation *in vivo* ^(37,38)^.

In addition, the Piezo1 agonist Yoda2 enhances the bone formation response to loading in aged mice ^(39)^. A role for Piezo1 and YAP/TAZ modulation in the Wnt1-mediated synergistic induction by loading and Scl/Dkk1 antibody warrants further investigation.

### Scl/Dkk1 antibody treatment, independent of loading, robustly increased endosteal bone formation in aged mice

We observed a highly significant main effect of antibody treatment on osteogenic gene expression, cortical and trabecular bone morphology, and dynamic histomorphometry outcomes, indicating that short-term dual antibody treatment increased bone formation and bone mass in aged mice. This is consistent with prior studies that demonstrated short-term (2-8 weeks) targeting of Scl + Dkk1 increased bone formation in 20-month old mice ^(40)^ and in 11-month old rats with OVX-induced osteoporosis ^(13)^. In particular, we saw that antibody treatment potently induced bone formation on the endocortical surface by increasing both mineralizing surface (approx. 5-fold) and mineral apposition rate (approx. 4-fold) (Fig. 3D, non-loaded tibias). By comparison, while bone formation on the periosteal surface also increased due to large fold-change increases in MS/BS and MAR (Fig. 3B), absolute values of periosteal indices were substantially lower compared to the endocortical surface (e.g., mean Ps.BFR/BS = 0.18 μm^3^/μm^2^/day, Ec.BFR/BS = 2.08; Fig. 3). Choi et al. ^(40)^ similarly reported that dual-antibody therapy of 20-month-old mice induced approximately 3-fold greater rate of bone formation on the endocortical compared to the periosteal surface of femurs. Consistent with the greater endosteal response, we observed a 50% increase in tibial trabecular bone volume compared to a 24% increase in cortical bone area (Figure S2A). (It should be noted that the proximal tibia in 22-month C57Bl/6 mice has very low bone volume (BV/TV ∼0.01), so even a 50% increase corresponds to a small absolute effect.) Interestingly, Florio et al. ^(13)^ reported that dual antibody treatment in 10-wk old mice and 11-mo old rats induced potent diaphyseal bone formation responses on both periosteal and endocortical surfaces. Taken together, while periosteal responses vary from modest to strong in studies using rodents of different ages, the endocortical/endosteal response is consistently strong. Thus, dual Scl/Dkk1 antibody therapy can potently improve bone formation to increase cortical thickness and trabecular bone volume, which are diminished in osteoporotic patients.

Our study is not without limitations. First, our experimental design did not include monotherapy targeting of Scl or Dkk1, which would allow for characterization of the independent effects of each antibody compared to their combined effect. Previous work has demonstrated that combination antibody therapy can be more potent than monotherapy, when evaluated without loading. For example, Choi et al. ^(40)^ reported that in both adult and aged mice, combination Scl-Ab + Dkk1-Ab at a total dose of 12.5 mg/kg had similar skeletal benefits as Scl-Ab alone at 25 mg/kg, suggesting that low-dose combination therapy can be as effective as a higher dose of Scl-Ab monotherapy. These findings suggest that the bone anabolic effects of combination therapy in our study (25 mg/kg total dose) might be superior to (or at least not less than) the effects of a similar total dose of Scl-Ab alone. (Note that Dkk1-Ab monotherapy has negligible skeletal benefits in normal mice; Dkk1-Ab is only effective when combined with Scl-Ab or when *Sost* is deleted ^(16,41)^.) However, the previous reports did not include a loading variable, and our study only tested dual antibody therapy at a single dose, therefore we cannot conclude that our antibody treatment would be superior to an equivalent dose of Scl-Ab alone in enhancing the effects of mechanical loading. A second limitation is that we used only a single loading magnitude (peak strain -2200 µ). This magnitude induced modest increases in bone formation (Fig. 3), but no increases in bone morphology based on serial microCT (Fig. 2). It is expected that loading to a higher strain value would have induced a greater anabolic effect, which may have resulted in detectable changes in bone morphology and perhaps revealed further synergistic effects. It is also likely that a longer duration study would result in more bone apposition due to loading, which would be detectable by serial microCT. Third, we included only female mice in our study. Few studies have directly compared the anabolic effects of Scl-Ab treatment (without loading) in male and female mice, but those that did reported similar effects in wildtype mice of both sexes ^(42,43)^. Studies on the combined effects Scl-Ab plus mechanical loading have been limited to female mice ^(30,31)^. Yang et al. ^(44)^ reported that the response to mechanical loading in Sost-deficient mice was sex dependent, suggesting that future work on combination antibody plus loading treatment should be done in both sexes. Lastly, mice are a model system and results may not translate directly to humans.

In conclusion, we showed that dual Scl-Ab + Dkk1-Ab treatment can potently increase bone formation in aged mice, and synergize with mechanical loading to enhance periosteal bone formation. This synergistic effect may be linked to upregulation of *Wnt1* expression. These results confirm prior results on the potent anabolic effect of dual Scl/Dkk1 antibody treatment in preclinical models of osteoporosis and extend these findings by demonstrating their effects when combined with mechanical loading. Our results support the concept that combinatorial therapy using dual Scl/Dkk1 antibodies and weight-bearing exercise may enhance bone formation in the osteoporotic skeleton and is a topic that warrants future clinical investigation.

## Acknowledgements

Funding support from the NIH: R01 AR047867, T32 GM007200, T32 AR060719, T32 EB018266, and F30 AG066310. MicroCT scanning performed at the Washington University Musculoskeletal Research, supported by P30 AR074992 and S10 RR023660. Sclerostin and Dkk1 antibodies provided as a gift from Amgen.

---

**Figure S1.**
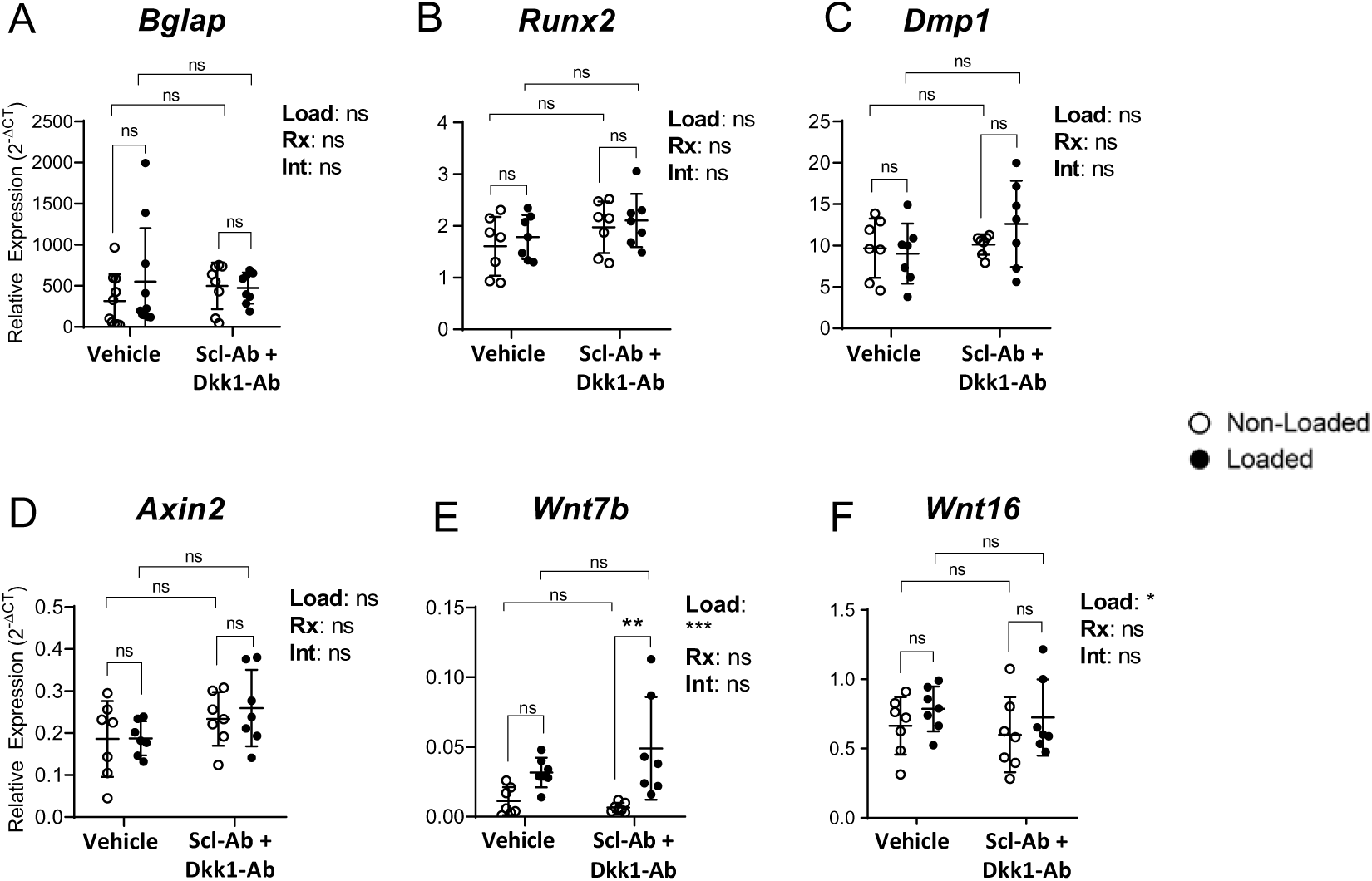
Osteogenic (A-C) and Wnt pathway-related (D-F) gene expression was surveyed by RT-qPCR after 5 consecutive days of loading (Fig. 1A). Treatment groups included vehicle control vs dual antibody (12.5 mg/kg Scl-Ab + 12.5 mg/kg Dkk1-Ab). The main effects of loading (“Load”), antibody treatment (“Rx”), and their interaction (“Int”) was evaluated by 2-factor repeated measures ANOVA. Sidak’s multiple comparisons *post hoc* test was used to compare differences between groups. \**p*<0.05; \*\**p*<0.01, \*\*\**p*<0.001, \*\*\*\**p*<0.0001; n=7-10/group.

**Figure S2.**
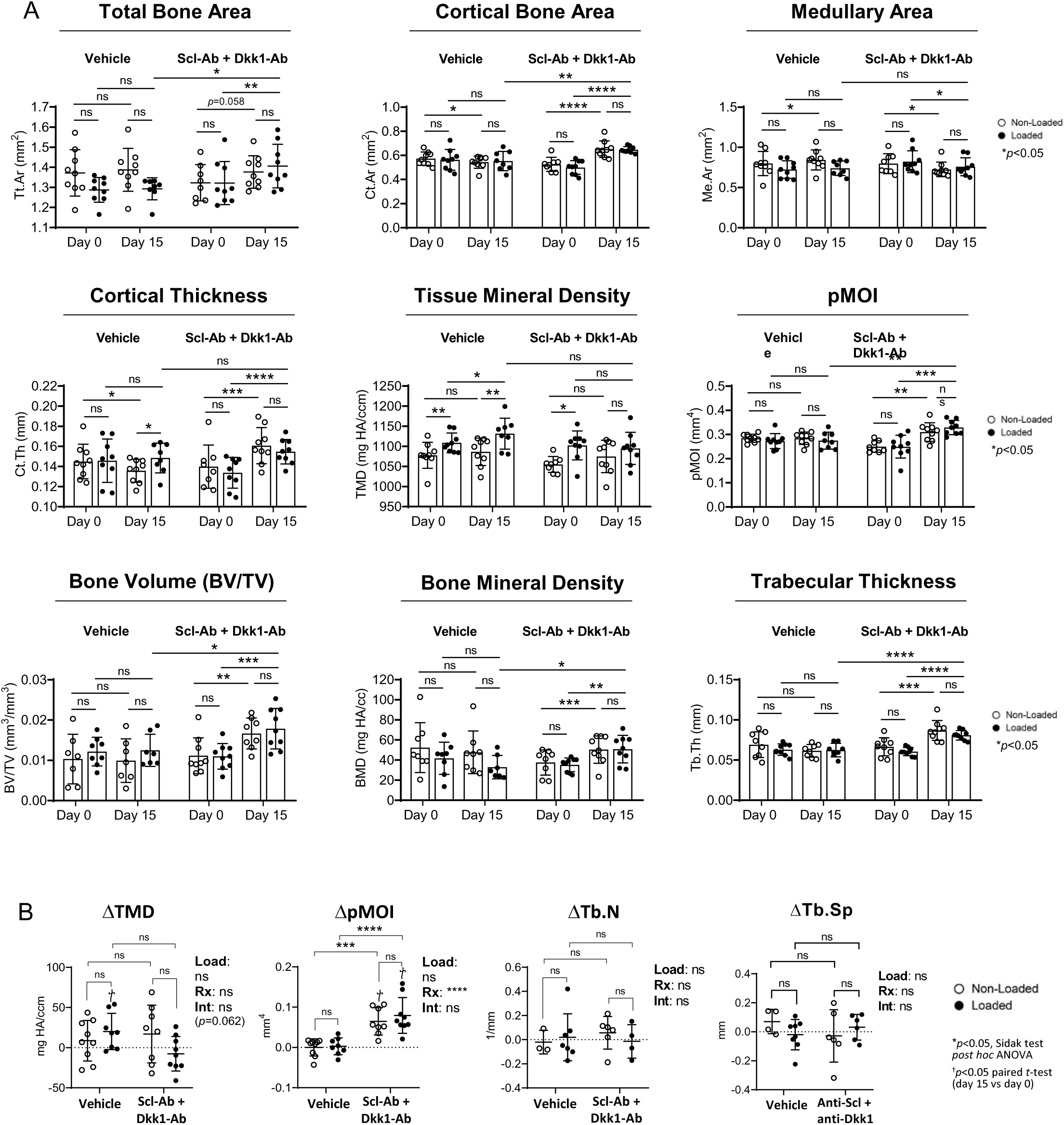
Serial µCT analysis. Bilateral tibias were serially scanned to evaluate the effects of loading and dual antibody treatment on bone morphology in 22-mo old mice. (A) Cortical (Tt.Ar, Ct.Ar, Me.Ar, Ct.Th, TMD, and pMOI) and trabecular (BV/TV, BMD, and Tb.Th) bone outcomes on days 0 and 15. Paired *t*-tests were used to determine if bone morphology was different on day 0 vs day 15. (B) Morphometric changes (Δ) in the loaded and contralateral non-loaded limbs of vehicle and antibody-treated mice. Main effects were evaluated by 2-factor repeated measures ANOVA, followed by Sidak’s multiple comparisons test. \**p*<0.05, \*\**p*<0.01, \*\*\**p*<0.001, and \*\*\*\**p*<0.0001; n=8-9/group. In (B) D indicates that there was a significant change (*p*<0.05, paired *t*-test) in morphology between days 0 and 15 (i.e., value different from zero; also shown in Fig S2A).

## Notes

**Grant Support:** This work was supported by NIH grants R01-AR047867 [MJS], P30-AR074992 [MJS], T32-GM007200 [CCS], T32-AR060719[LYL], T32 EB018266 [NM], and F30-AG066310 [CCS].

**Conflict of Interest:** The authors declare there is no conflict of interest.

### Competing Interest Statement

The authors have declared no competing interest.

## References

1. Canalis E. Wnt signalling in osteoporosis: mechanisms and novel therapeutic approaches. Nature reviews Endocrinology. 2013;9(10):575–83.

2. Baron R, Kneissel M. WNT signaling in bone homeostasis and disease: from human mutations to treatments. Nat Med. 2013;19(2):179–92.

3. Ramchand SK, Seeman E. Advances and Unmet Needs in the Therapeutics of Bone Fragility. Frontiers in endocrinology. 2018;9:505.

4. van Bezooijen RL, ten Dijke P, Papapoulos SE, Lowik CW. SOST/sclerostin, an osteocyte-derived negative regulator of bone formation. Cytokine & growth factor reviews. 2005;16(3):319–27.

5. Poole KE, van Bezooijen RL, Loveridge N, Hamersma H, Papapoulos SE, Lowik CW, Reeve J. Sclerostin is a delayed secreted product of osteocytes that inhibits bone formation. FASEB J. 2005;19(13):1842–4.

6. Semenov M, Tamai K, He X. SOST is a ligand for LRP5/LRP6 and a Wnt signaling inhibitor. J Biol Chem. 2005;280(29):26770–5.

7. Ai M, Holmen SL, Van Hul W, Williams BO, Warman ML. Reduced affinity to and inhibition by DKK1 form a common mechanism by which high bone mass-associated missense mutations in LRP5 affect canonical Wnt signaling. Mol Cell Biol. 2005;25(12):4946–55.

8. Li X, Ominsky MS, Warmington KS, Morony S, Gong J, Cao J, Gao Y, et al. Sclerostin antibody treatment increases bone formation, bone mass, and bone strength in a rat model of postmenopausal osteoporosis. J Bone Miner Res. 2009;24(4):578–88.

9. Kedlaya R, Veera S, Horan DJ, Moss RE, Ayturk UM, Jacobsen CM, Bowen ME, et al. Sclerostin inhibition reverses skeletal fragility in an Lrp5-deficient mouse model of OPPG syndrome. Science translational medicine. 2013;5(211):211ra158.

10. Saag KG, Petersen J, Brandi ML, Karaplis AC, Lorentzon M, Thomas T, Maddox J, et al. Romosozumab or Alendronate for Fracture Prevention in Women with Osteoporosis. N Engl J Med. 2017;377(15):1417–27.

11. Cosman F, Crittenden DB, Adachi JD, Binkley N, Czerwinski E, Ferrari S, Hofbauer LC, et al. Romosozumab Treatment in Postmenopausal Women with Osteoporosis. N Engl J Med. 2016;375(16):1532–43.

12. Taylor S, Ominsky MS, Hu R, Pacheco E, He YD, Brown DL, Aguirre JI, et al. Time-dependent cellular and transcriptional changes in the osteoblast lineage associated with sclerostin antibody treatment in ovariectomized rats. Bone. 2016;84:148–59.

13. Florio M, Gunasekaran K, Stolina M, Li X, Liu L, Tipton B, Salimi-Moosavi H, et al. A bispecific antibody targeting sclerostin and DKK-1 promotes bone mass accrual and fracture repair. Nat Commun. 2016;7:11505.

14. Holdsworth G, Greenslade K, Jose J, Stencel Z, Kirby H, Moore A, Ke HZ, et al. Dampening of the bone formation response following repeat dosing with sclerostin antibody in mice is associated with up-regulation of Wnt antagonists. Bone. 2018;107:93–103.

15. Li X, Grisanti M, Fan W, Asuncion FJ, Tan HL, Dwyer D, Han CY, et al. Dickkopf-1 regulates bone formation in young growing rodents and upon traumatic injury. J Bone Miner Res. 2011;26(11):2610–21.

16. Witcher PC, Miner SE, Horan DJ, Bullock WA, Lim KE, Kang KS, Adaniya AL, et al. Sclerostin neutralization unleashes the osteoanabolic effects of Dkk1 inhibition. JCI Insight. 2018;3(11).

17. Florio M, Kostenuik PJ, Stolina M, Asuncion FJ, Grisanti M, Ke HZ, Ominsky MS. Dual Inhibition of the Wnt Inhibitors DKK1 and Sclerostin Promotes Fracture Healing and Increases the Density and Strength of Uninjured Bone: An Experimental Study in Nonhuman Primates. J Bone Joint Surg Am. 2023;105(15):1145–55.

18. Angitia-Biopharmaceuticals. A First-in-Human Study Evaluating AGA2118 in Men and Postmenopausal Women. 2024.

19. Sawakami K, Robling AG, Ai M, Pitner ND, Liu D, Warden SJ, Li J, et al. The Wnt co-receptor LRP5 is essential for skeletal mechanotransduction but not for the anabolic bone response to parathyroid hormone treatment. J Biol Chem. 2006;281(33):23698–711.

20. Robinson JA, Chatterjee-Kishore M, Yaworsky PJ, Cullen DM, Zhao W, Li C, Kharode Y, et al. Wnt/beta-catenin signaling is a normal physiological response to mechanical loading in bone. J Biol Chem. 2006;281(42):31720–8.

21. Choi RB, Robling AG. The Wnt pathway: An important control mechanism in bone’s response to mechanical loading. Bone. 2021;153:116087.

22. Robling AG, Niziolek PJ, Baldridge LA, Condon KW, Allen MR, Alam I, Mantila SM, et al. Mechanical stimulation of bone in vivo reduces osteocyte expression of Sost/sclerostin. J Biol Chem. 2008;283(9):5866–75.

23. Holguin N, Brodt MD, Silva MJ. Activation of Wnt Signaling by Mechanical Loading Is Impaired in the Bone of Old Mice. J Bone Miner Res. 2016;31(12):2215–26.

24. Tu X, Rhee Y, Condon KW, Bivi N, Allen MR, Dwyer D, Stolina M, et al. Sost downregulation and local Wnt signaling are required for the osteogenic response to mechanical loading. Bone. 2012;50(1):209–17.

25. Kelly NH, Schimenti JC, Ross FP, van der Meulen MC. Transcriptional profiling of cortical versus cancellous bone from mechanically-loaded murine tibiae reveals differential gene expression. Bone. 2016;86:22–9.

26. Lawson LY, Brodt MD, Migotsky N, Chermside-Scabbo CJ, Palaniappan R, Silva MJ. Osteoblast-Specific Wnt Secretion Is Required for Skeletal Homeostasis and Loading-Induced Bone Formation in Adult Mice. J Bone Miner Res. 2022;37(1):108–20.

27. Lawson LY, Migotsky N, Chermside-Scabbo CJ, Shuster JT, Civitelli R, Silva MJ. Loading-Induced Bone Formation is Mediated by Wnt1 Induction in Osteoblast-Lineage Cells. bioRxiv. 2022.

28. Wergedal JE, Kesavan C, Brommage R, Das S, Mohan S. Role of WNT16 in the regulation of periosteal bone formation in female mice. Endocrinology. 2015;156(3):1023–32.

29. Chermside-Scabbo CJ, Harris TL, Brodt MD, Braenne I, Zhang B, Farber CR, Silva MJ. Old Mice Have Less Transcriptional Activation But Similar Periosteal Cell Proliferation Compared to Young-Adult Mice in Response to in vivo Mechanical Loading. J Bone Miner Res. 2020;35(9):1751–64.

30. Morse A, Schindeler A, McDonald MM, Kneissel M, Kramer I, Little DG. Sclerostin Antibody Augments the Anabolic Bone Formation Response in a Mouse Model of Mechanical Tibial Loading. J Bone Miner Res. 2018;33(3):486–98.

31. Gerbaix M, Ammann P, Ferrari S. Mechanically Driven Counter-Regulation of Cortical Bone Formation in Response to Sclerostin-Neutralizing Antibodies. J Bone Miner Res. 2021;36(2):385–99.

32. Main RP, Shefelbine SJ, Meakin LB, Silva MJ, van der Meulen MCH, Willie BM. Murine Axial Compression Tibial Loading Model to Study Bone Mechanobiology: Implementing the Model and Reporting Results. J Orthop Res. 2020;38(2):233–52.

33. Patel TK, Brodt MD, Silva MJ. Experimental and finite element analysis of strains induced by axial tibial compression in young-adult and old female C57Bl/6 mice. J Biomech. 2014;47(2):451–7.

34. Bouxsein ML, Boyd SK, Christiansen BA, Guldberg RE, Jepsen KJ, Muller R. Guidelines for assessment of bone microstructure in rodents using micro-computed tomography. J Bone Miner Res. 2010;25(7):1468–86.

35. Dempster DW, Compston JE, Drezner MK, Glorieux FH, Kanis JA, Malluche H, Meunier PJ, et al. Standardized nomenclature, symbols, and units for bone histomorphometry: a 2012 update of the report of the ASBMR Histomorphometry Nomenclature Committee. J Bone Miner Res. 2013;28(1):2–17.

36. Holguin N, Silva MJ. In-Vivo Nucleus Pulposus-Specific Regulation of Adult Murine Intervertebral Disc Degeneration via Wnt/Beta-Catenin Signaling. Sci Rep. 2018;8(1):11191.

37. Li X, Han L, Nookaew I, Mannen E, Silva MJ, Almeida M, Xiong J. Stimulation of Piezo1 by mechanical signals promotes bone anabolism. Elife. 2019;8.

38. Sasaki F, Hayashi M, Mouri Y, Nakamura S, Adachi T, Nakashima T. Mechanotransduction via the Piezo1-Akt pathway underlies Sost suppression in osteocytes. Biochemical and biophysical research communications. 2020;521(3):806–13.

39. Meslier QA, Oehrlein R, Shefelbine SJ. Combined Effects of Mechanical Loading and Piezo1 Chemical Activation on 22-Months-Old Female Mouse Bone Adaptation. Aging Cell. 2025;24(8):e70087.

40. Choi RB, Hoggatt AM, Horan DJ, Rogers EZ, Hong JM, Robling AG. Targeting Sclerostin and Dkk1 at Optimized Proportions of Low-Dose Antibody Achieves Similar Skeletal Benefits to Higher-Dose Sclerostin Targeting in the Mature Adult and Aged Skeleton. Aging Dis. 2022;13(6):1891–900.

41. Choi RB, Bullock WA, Hoggatt AM, Loots GG, Genetos DC, Robling AG. Improving Bone Health by Optimizing the Anabolic Action of Wnt Inhibitor Multitargeting. JBMR Plus. 2021;5(5):e10462.

42. Cardinal M, Chretien A, Roels T, Lafont S, Ominsky MS, Devogelaer JP, Manicourt DH, et al. Gender-Related Impact of Sclerostin Antibody on Bone in the Osteogenesis Imperfecta Mouse. Front Genet. 2021;12:705505.

43. Carpenter KA, Ross RD. Sclerostin Antibody Treatment Increases Bone Mass and Normalizes Circulating Phosphate Levels in Growing Hyp Mice. J Bone Miner Res. 2020;35(3):596–607.

44. Yang H, Buttner A, Albiol L, Julien C, Thiele T, Figge C, Kramer I, et al. Cortical bone adaptation to a moderate level of mechanical loading in male Sost deficient mice. Sci Rep. 2020;10(1):22299.

